# Characterizing Inpatient Medicine Resident Electronic Health Record Usage Patterns Using Event Log Data

**DOI:** 10.1101/428169

**Authors:** Jonathan H. Chen, Jason K. Wang, David Ouyang, Jason Hom, Jeffrey Chi

## Abstract

Amid growing rates of burnout, physicians report increasing electronic health record (EHR) usage alongside decreasing clinical facetime with patients. There exists a pressing need to improve physician-computer-patient interactions by streamlining EHR workflow.

To identify interventions to improve EHR design and usage, we systematically characterize EHR activity among internal medicine residents at a tertiary academic hospital across various inpatient rotations and roles from June 2013 to November 2016.

Logged EHR timestamps were extracted from Stanford Hospital’s EHR system (Epic) and cross-referenced against resident rotation schedules. We tracked the quantity of EHR logs across 24-hour cycles to reveal daily usage patterns. In addition, we decomposed daily EHR time into time spent on specific EHR actions (e.g. chart review, note entry and review, results review).

In examining 24-hour usage cycles from general medicine day and night team rotations, we identified a prominent trend in which night team activity promptly ceased at the shift’s end, while day team activity tended to linger post-shift. Across all rotations and roles, residents spent on average 5.38 hours (standard deviation=2.07) using the EHR. PGY1 (post-graduate year one) interns and PGY2+ residents spent on average 2.4 and 4.1 times the number of EHR hours on information review (chart, note, and results review) as information entry (note and order entry).

Analysis of EHR event log data can enable medical educators and programs to develop more targeted interventions to improve physician-computer-patient interactions, centered on specific EHR actions.

## INTRODUCTION

As medical educators, we hope our resident trainees value direct patient care and contact. Instead, we may progressively find their attention dominated by electronic health records (EHR) that mediate their work and proxy for their patients.^1^ Observational studies confirm an increasing shift from direct patient care to computer use in the wake of duty hour restrictions,^2,3,4^ informing ongoing reforms into the structure of medical training.^5^ Physicians report increasing time spent on paperwork and the computer^6^, alongside less time available for clinical interactions with patients.^7^ A growing burden of redundant clinical notes, alert fatigue, and an overflowing inbox has led to a systemic “4000 clicks a day” problem^8^ that has contributed to physician job dissatisfaction and burnout rates. Indeed, correlations between an increasing EHR task load and physician burnout have been demonstrated.^6,9^ There exists a pressing need to understand clinical EHR usage to inform opportunities to improve effective patient care processes.

Previous studies have quantified EHR usage intensity through direct observation and self-reported diaries.^2,10^ Ironically, with the amount of time trainees spend on the EHR, the computer also provides precise, reproducible, and scalable quantification of their electronic activities. Prior comparisons between direct observation of clinician activity and logged EHR timestamps confirm that the two approaches yield similar results for workflow analysis.^11,12^ Here, we use timestamp data to systematically characterize the intensity of EHR usage across different inpatient medicine rotations and roles.

## METHODS

Logged timestamps from the Epic EHR system for inpatient rotations of internal medicine residents at an academic tertiary care hospital were extracted from the STRIDE project^13^ from June 2013 to November 2016. We tracked the quantity of logged EHR actions accessed over 24-hour cycles per user binning by half-hour intervals. Logged EHR actions correspond to behaviors performed on the EHR as clinicians navigate components (e.g. notes, orders, results) of a patient’s electronic chart. To estimate time spent per action, we considered time intervals between access logs separated by 5 minutes or less of inactivity. Sensitivity of results relative to time intervals up to 10 minutes was also considered. We binned EHR actions into common behavioral categories and computed mean time spent per category per work day. Daily EHR usage was estimated as the sum of all active inter-access time intervals per day. User timestamps were cross-referenced against resident year and rotation schedules to account for each user’s progression through the internal medicine residency program over the three-year window. User-days with less than one hour of activity were excluded from analysis to account for remote access during vacation days. P-values were computed using two-sample t-tests allowing unequal variances (Welch’s t-test). Analyses were performed with Python 2.7 and R 2.13. The study was deemed non-human subjects research by the Stanford Institutional Review Board.

## RESULTS

During the three-year window, 15,909,629 unique actions were logged by 101 unique residents covering 99 PGY1 (post-graduate year one) intern-years and 61 PGY2+ resident-years. Fig 1 illustrates the intensity of EHR interactions per resident day during a 24-hour cycle for PGY1 interns versus PGY2+ residents across different inpatient rotations.

**Fig 1:**
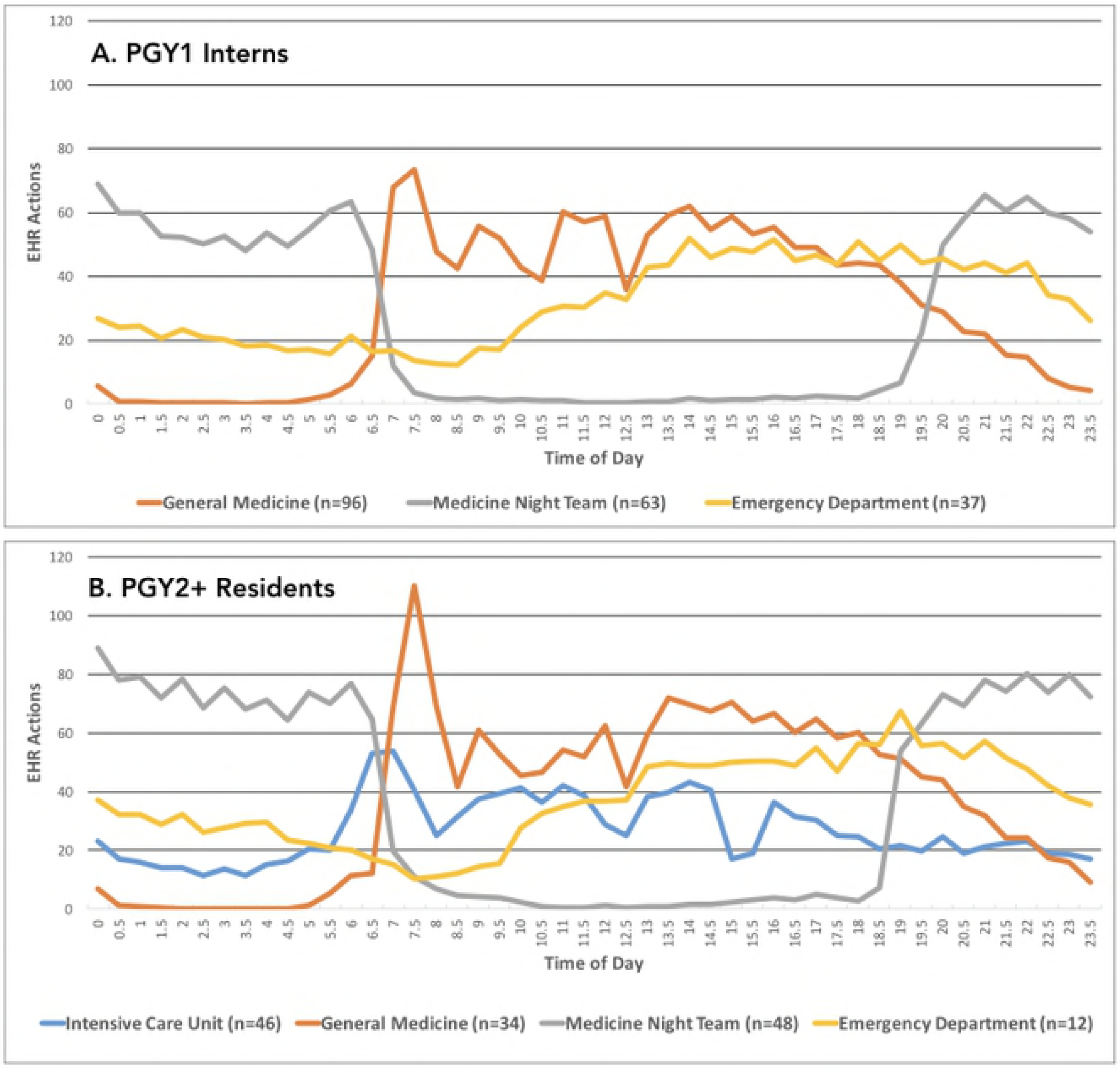
EHR Actions Vs. Time of Day

Table 1 highlights mean daily EHR usage times, decomposed by mean time spent on common EHR action categories across each inpatient rotation and role. Across all rotations, trainees spent a mean of 5.38 hours (PGY1=5.47, PGY2+=5.21) with standard deviation=2.07 (PGY1=2.10, PGY2+=2.02) and median of 5.33 hours (PGY1=5.47, PGY2+=5.09) with interquartile range=3.85-6.85 (PGY1=3.90-6.97, PGY2+=3.80-6.65) per work day using the EHR. Medical chart reviews account for the largest portion of activity (>40% on average) followed primarily by note entry and review. PGY1 interns spent disproportionately more time on note entry compared to PGY2+ residents during general medicine (15.2% vs. 4.0% of daily EHR usage, P<0.001) and emergency medicine rotations (18.4% vs. 13.9%, P<0.001), but similar times during the night team rotation (11.9% vs. 11.3%, P=0.28). Conversely, across all three rotations, PGY2+ residents spent disproportionately more time on note review compared to PGY1 interns (general medicine=20.7% vs. 13.0%, P<0.001; night team=21.8% vs. 11.6%, P<0.001; emergency medicine=11.6% vs. 8.1%, P<0.001). Trainees spent roughly 7.6 minutes (PGY2+ only), 9.7 minutes (PGY1=12.3, PGY2+=6.6), 3.7 minutes (PGY1=3.5, PGY2+=4.2), and 7.2 minutes (PGY1=8.0, PGY2+=5.5) on chart review actions per patient across intensive care unit, general medicine, night team, and emergency department rotations, respectively.

**Table 1:** EHR Usage Intensity Across Inpatient Medical Rotations and Roles Per User-Day^a^. EHR usage summary statistics and mean time spent on common EHR action categories per user-day across different inpatient rotations for PGY1 interns and PGY2+ residents. Time spent on category-specific actions are also shown as a percentage of mean daily EHR usage time. The EHR Navigator consists of pre-curated sequences of modules to facilitate common actions such as admission, rounding, and discharge.

| PGY1 Interns | General<br>Medicine<br>(n=96) | Medicine<br>Night Team<br>(n=63) | Emergency<br>Department<br>(n=37) |
| --- | --- | --- | --- |
| Median EHR Actions (IQR) | 1669 (1224-2167) | 1240 (901-1709) | 1704 (1161-2286) |
| Median % EHR Actions Accessed Remotely (IQR)<sup>b</sup> | 1% (0-8%) | 0% (0-7%) | 0% (0-3%) |
| Median Patient Records Accessed (IQR) | 11 (8-16) | 33 (24-44) | 21 (14-31) |
| Mean Time Spent Per EHR Action Category (Hours, % Daily) |  |  |  |
| Chart Review | 2.26 (40.4%) | 1.95 (42.9%) | 2.80 (46.1%) |
| Note Review | 0.73 (13.0%) | 0.53 (11.6%) | 0.49 (8.1%) |
| Note Entry | 0.85 (15.2%) | 0.54 (11.9%) | 1.12 (18.4%) |
| Order Entry | 0.64 (11.4%) | 0.50 (11.0%) | 0.38 (6.3%) |
| Navigator | 0.35 (6.3%) | 0.37 (8.1%) | 0.44 (7.2%) |
| Results Review | 0.26 (4.6%) | 0.35 (7.7%) | 0.28 (4.6%) |
| Mean Daily EHR Usage (SD) | 5.60 (2.03) | 4.55 (1.80) | 6.08 (2.40) |

**Table 1.** EHR usage summary statistics and mean time spent on common EHR action categories per user-day across different inpatient rotations for PGY1 interns and PGY2+ residents.
| PGY2+ Residents | General<br>Medicine<br>(n=34) | Medicine<br>Night Team<br>(n=48) | Emergency<br>Department<br>(n=12) | Intensive<br>Care Unit<br>(n=46) |
| --- | --- | --- | --- | --- |
| Median EHR Actions<br>(IQR) | 2008 (1340-2673) | 1741 (1266-2427) | 2125 (1317-2850) | 1427 (1105-1850) |
| Median % EHR Actions<br>Accessed Remotely<br>(IQR)<sup>b</sup> | 4% (0-13%) | 1% (0-7%) | 0% (0-2%) | 0% (0-5%) |
| Median Patient Records<br>Accessed (IQR) | 21 (16-27) | 31 (21-46) | 31 (20-45) | 17 (14-22) |
| Mean Time Spent Per<br>EHR Action Category<br>(Hours, % Daily) |  |  |  |  |
| Chart Review | 2.31 (43.9%) | 2.18 (39.6%) | 2.85 (46.5%) | 2.15 (45.9%) |
| Note Review | 1.09 (20.7%) | 1.20 (21.8%) | 0.71 (11.6%) | 0.75 (16.0%) |
| Note Entry | 0.21 (4.0%) | 0.62 (11.3%) | 0.85 (13.9%) | 0.20 (4.3%) |
| Order Entry | 0.50 (9.5%) | 0.54 (9.8%) | 0.38 (6.2%) | 0.44 (9.4%) |
| Navigator | 0.31 (5.9%) | 0.23 (4.2%) | 0.46 (7.5%) | 0.41 (8.8%) |
| Results Review | 0.27 (5.1%) | 0.25 (4.5%) | 0.25 (4.1%) | 0.29 (6.2%) |
| Mean Daily EHR Usage<br>(SD) | 5.26 (2.00) | 5.50 (2.17) | 6.13 (2.58) | 4.68 (1.56) |

<sup>a</sup>IQR = interquartile range, SD = standard deviation
<sup>b</sup>The EHR system is accessed either remotely (e.g. via mobile device) or via computer workstations situated in the hospital.

Daily EHR usage estimations prove robust to the choice of inactivity cutoff; the standard deviation of mean daily EHR usages computed using inactivity cutoffs from 5-10 minutes is 28.8 minutes. The choice of a 5-minute cutoff captures the vast majority of active computer sessions (98.4% of inter-access time intervals), while excluding lengthy idle sessions. Excluding user-days based on an activity threshold of one hour removes 11.6% of all user-days (435/3756), corresponding to roughly one day off per week.

## DISCUSSION AND CONCLUSION

With increasing reliance on EHRs to mediate patient care, direct analysis of EHR audit logs provides a granular and objective way to characterize physician EHR usage. A key limitation of this approach is the estimation of idle time between access logs; overestimating idle times could overestimate EHR usage times. However, our sensitivity analysis showed that estimations were robust to cutoffs between 5-10 minutes. Additionally, although results are aggregated across multiple years, the reported EHR usage statistics are derived from a single institution.

The pattern of EHR activity over 24-hour cycles provides qualitative insights into resident behavior (Figure 1). The 2011 ACGME duty hour restrictions prompted the separation of the hospital’s General Medicine rotation into Day and Night Teams. These 24-hour cycles may suggest that night teams treat patient care as discrete shift work, with clinical activities promptly ceasing at 7 AM as noted by the steep drop-off, while general medicine (day team) EHR activity often lingers well beyond duty hour recommendations (9 PM onwards) and restrictions (11 PM onwards), defined as ten and eight hours before the subsequent 7 AM shift.

Perhaps more concerning is that the prevalence of physician-computer interaction almost certainly restricts direct physician-patient interaction. Assuming a 12-hour work day^3^, PGY1 interns and PGY2+ residents spent nearly half (PGY1=46%, PGY2+=43%) of work time on the EHR, respectively. This is especially notable with peaked EHR activity when Day Teams arrive, suggesting the traditional model of pre-rounding at the patient bedside has been replaced by the workroom computer as the trusted source for patient information.^1^

Comparing between roles (Table 1), we see that PGY2+ residents execute a greater number of EHR actions with disproportionately more time spent on note review, yet less daily EHR time, compared to PGY1 interns. Indeed, in the clinic, PGY2+ residents embrace a supervisory role and often oversee twice the number of patients (note the nearly 2:1 median patient ratio during general medicine rotations) compared to PGY1 interns. Conversely, PGY1 interns are delegated time-intensive note entry duties albeit for a smaller population of patients, potentially accounting for greater daily EHR time. Across diverse rotations, we see significant time spent on the same pattern of EHR activities. PGY1 interns and PGY2+ residents spent on average 2.4 and 4.1 times the number of EHR hours on information review (chart, note, and results review) as information entry (note and order entry) (Table 1), respectively. Improvements in EHR design could target the burden of data retrieval by encouraging more concise documentation and redesigned or automated content organization.^14^ By identifying specific EHR activities that consistently dominate resident computer usage across multiple inpatient rotations and roles, we hope to facilitate a more targeted, data-driven approach to improving physician-computer-patient interactions.

## ACKNOWLEDGEMENTS

Project supported by the Stanford Cyber Initiative. J.H.C was supported in part by the NIH Big Data 2 Knowledge Award Number K01ES026837 through the National Institute of Environmental Health Sciences.

Patient data extracted and de-identified by Gomathi Krishnan of the STRIDE (Stanford Translational Research Integrated Database Environment) project, a research and development project at Stanford University to create a standards-based informatics platform supporting clinical and translational research. The STRIDE project described was supported by the National Center for Research Resources and the National Center for Advancing Translational Sciences, National Institutes of Health, through grant UL1 RR025744.

Content is solely the responsibility of the authors and does not necessarily represent the official views of the NIH, or Stanford Healthcare.

## Contributions Statement

JHC conceived the study and design. JHC and JKW performed analyses and drafted the initial manuscript. All authors reviewed the manuscript and contributed to study design.

